# Repeated horizontal gene transfer of *GAL*actose metabolism genes violates Dollo’s law of irreversible loss

**DOI:** 10.1101/2020.07.22.216101

**Authors:** Max A. B. Haase, Jacek Kominek, Dana A. Opulente, Xing-Xing Shen, Abigail L. LaBella, Xiaofan Zhou, Jeremy DeVirgilio, Amanda Beth Hulfachor, Cletus P. Kurtzman, Antonis Rokas, Chris Todd Hittinger

**Author notes:** Deceased (Cletus P. Kurtzman). Equal authorship (Max A. B. Haase, Jacek Kominek).

## Abstract

Dollo’s law posits that evolutionary losses are irreversible, thereby narrowing the potential paths of evolutionary change. While phenotypic reversals to ancestral states have been observed, little is known about their underlying genetic causes. The genomes of budding yeasts have been shaped by extensive reductive evolution, such as reduced genome sizes and the losses of metabolic capabilities. However, the extent and mechanisms of trait reacquisition after gene loss in yeasts have not been thoroughly studied. Here, through phylogenomic analyses, we reconstructed the evolutionary history of the yeast galactose utilization pathway and observed widespread and repeated losses of the ability to utilize galactose, which occurred concurrently with the losses of *GAL*actose (*GAL*) utilization genes. Unexpectedly, we detected three galactose-utilizing lineages that were deeply embedded within clades that underwent ancient losses of galactose utilization. We show that at least two, and possibly three, lineages reacquired the *GAL* pathway via yeast-to-yeast horizontal gene transfer. Our results show how trait reacquisition can occur tens of millions of years after an initial loss via horizontal gene transfer from distant relatives. These findings demonstrate that the losses of complex traits and even whole pathways are not always evolutionary dead-ends, highlighting how reversals to ancestral states can occur.

## Introduction

Understanding the interactions between a species’ phenotype, genotype, and environment is a central goal of evolutionary biology. Of particular interest are the mechanisms by which the environment selects for changes in phenotype and subsequently genome content. Due to their remarkable physiological diversity, budding yeasts are present in an extraordinary range of environments^1^. Alongside robustly characterized physiologies^2^ and the availability of an unrivaled set of genome sequences^1,3,4^, budding yeasts provide a unique subphylum-level eukaryotic model for studying the interplay between the genome, phenotype, and the environment.

Trait reversal is an intriguing phenomenon whereby the character state of a particular evolutionary lineage returns to its ancestral state. For more than a century, trait reversal after a loss event has been thought to be highly unlikely; Dollo’s law of irreversibility states that, once a trait is lost, it is unlikely for the same trait to be found in a descendant lineage, thereby excluding certain evolutionary paths^5,6^. Despite this purist interpretation, many examples of apparent violations to Dollo’s law have been documented^7–15^, and it is clear that evolutionary processes sometimes break Dollo’s law^16–18^. Nonetheless, the molecular and genetic mechanisms leading to trait reversal have only been determined in a few cases^17,18^. For example, it was recently shown that flower color reversal in a *Petunia* species was facilitated by the resurrection of a pseudogene^18^. In this case, the reversal was temporally rapid, which is in agreement with the hypothesis that traits flicker on and off during speciation^16^. These results underscore that complex traits do indeed undergo reversal and help identify one possible genetic mechanism for doing so. In other cases, traits have been reversed long after the speciation process and long after pseudogenes are undetectable^7,19^, raising the question of how trait reversal can occur millions of years after the initial loss.

The Leloir pathway of galactose utilization in the model budding yeast *Saccharomyces cerevisiae* (subphylum Saccharomycotina) is one of the most intensely studied and well-understood genetic, regulatory, and metabolic pathways of any eukaryote^20–29^. Although its regulatory genes are unlinked, the *GAL* genes encoding the three key catabolic enzymes (*GAL1, GAL7*, and *GAL10*) are present in a localized gene cluster^25^. A critical consequence of clustering genes in fungi is a marked increase in the rate of gene loss^22,25,30–32^ and a striking increase in the incidence of horizontal gene transfer (HGT) of those genes^32,33^. The principal mode of evolution for the *GAL* gene cluster has been differential gene loss from an ancestral species that possessed the *GAL* genes in a cluster^4,22,25,34^. In one case, the budding yeast *GAL* enzymatic gene cluster was horizontally transferred into the fission yeast *Schizosaccharomyces pombe* (subphylum Taphrinomycotina)^25^. Nonetheless, this transferred cluster is not functional in typical growth assays, suggesting *Sc. pombe GAL* cluster may not be deployed catabolically or may respond to induction signals other than galactose^35^. Dairy and some other strains of *Saccharomyces cerevisiae* may have horizontally acquired a more active, transcriptionally rewired *GAL* pathway from an unknown outgroup of the genus *Saccharomyces*^36,37^, or they may have preserved these two versions of the pathway through extreme balancing selection^38^, but trait reversal is highly unlikely under either interpretation. Collectively, these prior observations suggest that both cis-regulatory features and unlinked regulators play crucial roles in determining the function of horizontally transferred genes. Due to the widespread loss of *GAL* genes and the apparent ability for the *GAL* enzymatic gene cluster to be horizontally transferred intact, we hypothesized that budding yeast *GAL* clusters might break Dollo’s law under some conditions.

To address this hypothesis, we explored the genetic content and phenotypic capabilities of a diverse set of budding yeast genomes. Despite being deeply embedded within clades that underwent ancient losses of galactose metabolism, the genera *Brettanomyces* and *Wickerhamomyces* both contained representatives that could utilize galactose. Analyses of their genome sequences revealed *GAL* gene clusters that exhibited an unusually high degree of synteny with gene clusters in distantly related species. Further analysis of the genome of *Nadsonia fulvescens* showed that it also contains a *GAL* gene cluster that is remarkably similar to a distantly related species.

Through rigorous phylogenetic hypothesis testing, we found strong evidence for the complete losses of the genes encoding the enzymes necessary for galactose catabolism, followed by their reacquisitions via independent yeast-to-yeast HGT events in at least two, and possibly three, cases. Genes lost in budding yeasts have been regained via HGT from bacterial donors in several cases ^39–45^, but here we demonstrate an exceptionally clear example of a complex trait and its corresponding genes being lost and then regained to its ancestral eukaryotic form. We conclude that multiple distantly related lineages of yeasts have circumvented evolutionary irreversibility, both at the molecular and phenotypic level, via eukaryotic HGT and that evolutionary paths are not absolutely constrained after trait loss.

## Results

### Genome selection and sequencing

To reconstruct the evolution of galactose metabolism in the budding yeast subphylum Saccharomycotina, we first selected a set of genomes to analyze that spanned the backbone of the subphylum^3,4^. Next, we sequenced the genomes of five additional species at strategically positioned branches: *Brettanomyces naardenensis*; a yet-to-be described *Wickerhamomyces* species, *Wickerhamomyces* sp. UFMG-CM-Y6624; *Candida chilensis*; *Candida cylindracea*; and *Candida silvatica*. All strains used in this study can be found in Supplemental Table 1. Finally, we reconstructed a species-level phylogeny, analyzing the genome sequences of 96 Saccharomycotina and 10 outgroup species (Supplemental Figures 1 and 2).

**Figure 1.**
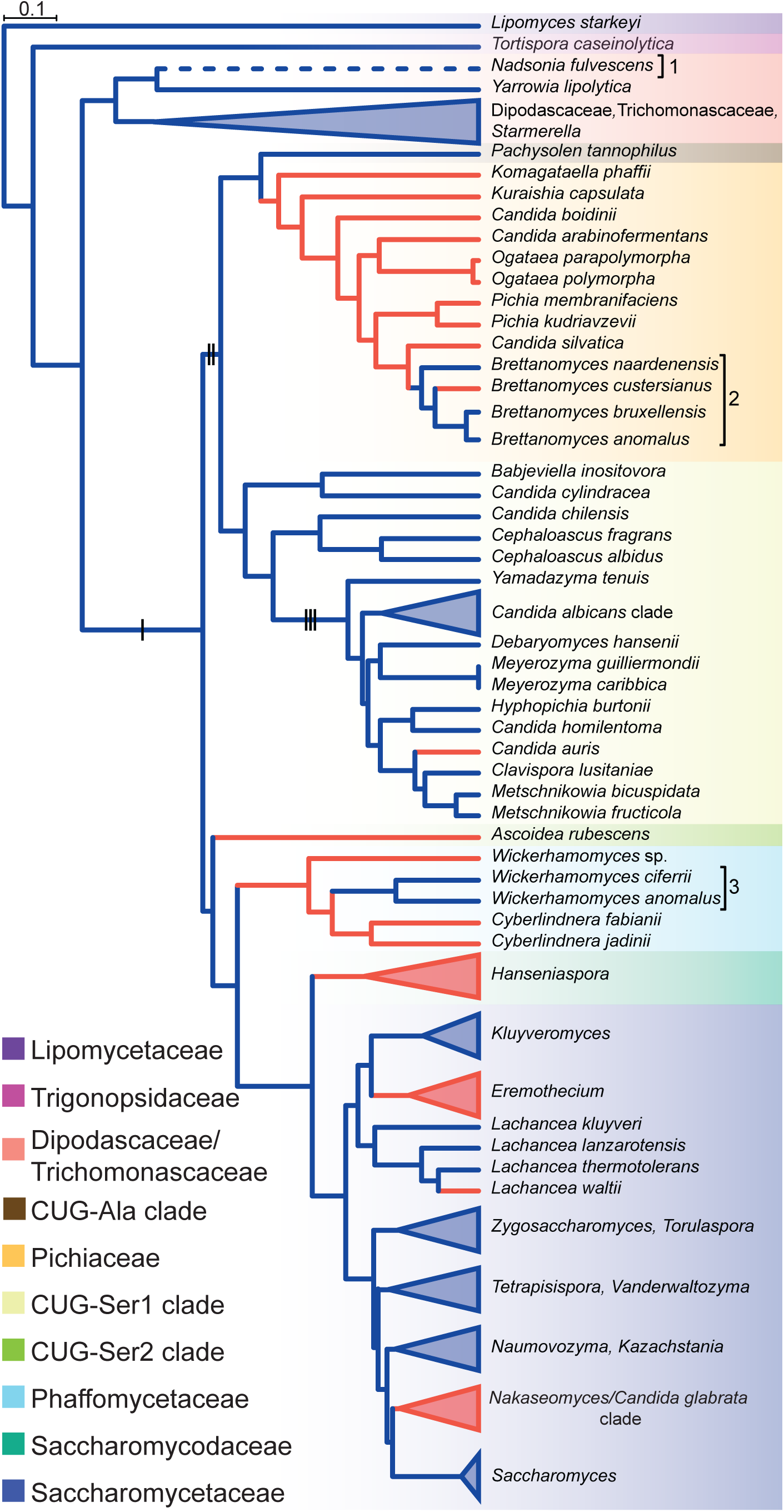
Evolutionary history of galactose metabolism in budding yeasts. Species-level presence or absence of galactose utilization is mapped onto the relative divergence timetree (Supplemental Figures 2 and 3) with some clades collapsed. Branch color denotes the ability to metabolize galactose; blue (+) and red(–). The black bars mark the branches of three key events in the evolution of the *GAL* cluster (I-cluster formation, II-translocation of *ORF-Y* into the cluster, and III-translocation of *ORF-X* into the cluster). Numbered groups indicate the three clades with unexpected *GAL* clusters. The dashed branch of the *Nadsonia* lineage indicates the ambiguity of the ancestral character state due to its extremely long branch (Supplemental Figures 2 and 3).

**Figure 2.**
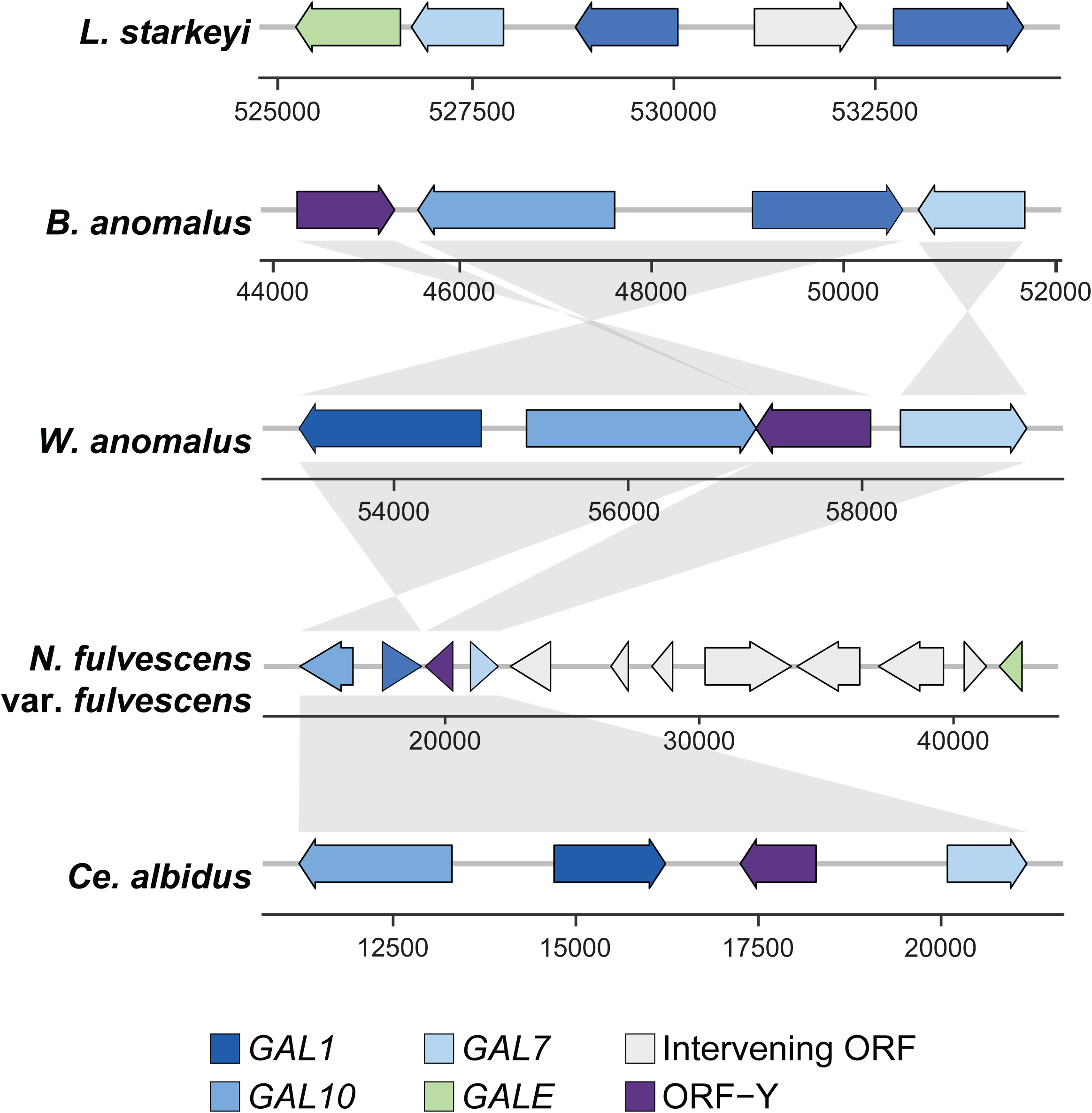
Surprisingly syntenic *GAL* clusters between distantly related groups of yeasts. The *GAL* clusters of five representative species are shown. Numbers correspond to positions in each scaffold or contig. Further details and examples are provided in Supplemental Figures 3 and 4.

### Recurrent loss of yeast *GAL* clusters

This dataset suggests that the *GAL* enzymatic gene cluster (hereafter *GAL* cluster) of budding yeasts formed prior to the last common ancestor of the CUG-Ser1, CUG-Ser2, CUG-Ala, Phaffomycetaceae, Saccharomycodaceae, and Saccharomycetaceae major clades (Figure 1 and Supplemental Figure 3)^25^. This inference is supported by the presence of the fused bifunctional *GAL10* gene in these lineages and the absence of the fused protein in species outside these lineages (Figure 1 and Supplemental Figure 3)^25^. Since galactose metabolism has been repeatedly lost over the course of budding yeast evolution and the enzymatic genes are present in a gene cluster, we next asked whether the trait of galactose utilization had undergone trait reversal. We reasoned that species or lineages who utilize galactose, but who are deeply embedded in clades that predominantly cannot utilize galactose, would represent prime candidates for possible trait reversal events. When we mapped both *GAL* gene presence and galactose utilization onto our phylogeny (Figure 1 and Supplemental Figure 3), we inferred repeated loss of the *GAL* gene clusters (Figure 1 and Supplemental Figure 3) and a strong association between genotype and phenotype (Supplemental Table 2). However, we identified two genera, *Brettanomyces* and *Wickerhamomyces*, as containing candidates for trait reversal (Figure 1). This unusual trait distribution led us to consider the possibility that the *GAL* clusters of these two lineages were not inherited vertically.

**Figure 3.**
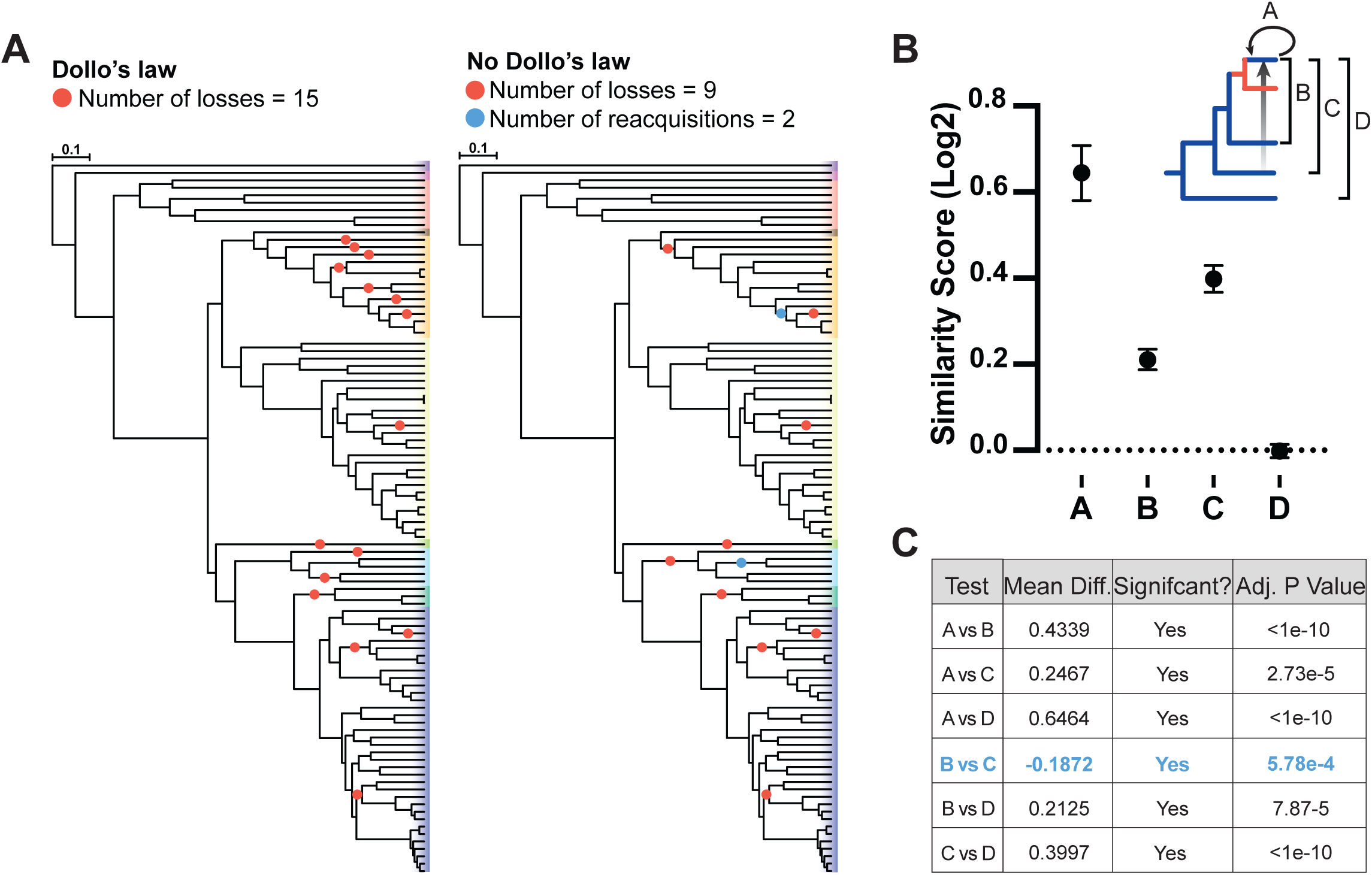
Comparison of Dollo’s law versus reacquisition of the *GAL* genes from the CUG-Ser1 clade. (**A**) Evolutionary trait reconstruction, based on a parsimony framework either assuming that traits cannot be regained (left) or that traits can be regained (right). (**B**) Similarity score of the Gal1, Gal7, and Gal10 proteins as calculated by protein sequence similarity and the comparisons shown in the upper right; means with standard deviations are depicted. Raw percent identity values are shown in Supplemental Figure 10. Comparisons used to calculate similarity scores: A, between species within the clade with potentially transferred *GAL* genes (recipient clade); B, between the recipient clade and their closest relative with *GAL* genes; C, between the recipient clade and the potential donor lineage (CUG-Ser1 clade); and D, between the recipient clade and an outgroup lineage. (**C**) Student’s t-test of the mean difference between groups. Negative values violate the assumptions of vertical inheritance, and the critical comparison between B and C is bolded in blue.

### Unusual synteny patterns of *GAL* clusters

If the observed distribution of galactose metabolism were to be explained by only vertical reductive evolution, then *GAL* cluster losses have occurred even more frequently than currently appreciated. Interestingly, we noted that the structures of *Brettanomyces* and *Wickerhamomyces GAL* clusters are strikingly syntenic to the *GAL* clusters belonging to distantly related yeasts, specifically those belonging to the CUG-Ser1 clade, which includes *Candida albicans* (Figure 2 and Supplemental Figure 4). Since the CUG-Ser1 clade *GAL* cluster structure is evolutionarily derived^25^, it is highly unlikely that these two additional lineages would independently evolve such similar structures. Instead, one might expect *Brettanomyces* to share a structure with *Pacchysolen tannophilus*, its closest relative containing a *GAL* cluster. These observations suggest, a model wherein the *Brettanomyces* and *Wickerhamomyces GAL* clusters share ancestry with *GAL* clusters from the CUG-Ser1 clade, rather than with those from their much closer organismal relatives.

**Figure 4.**
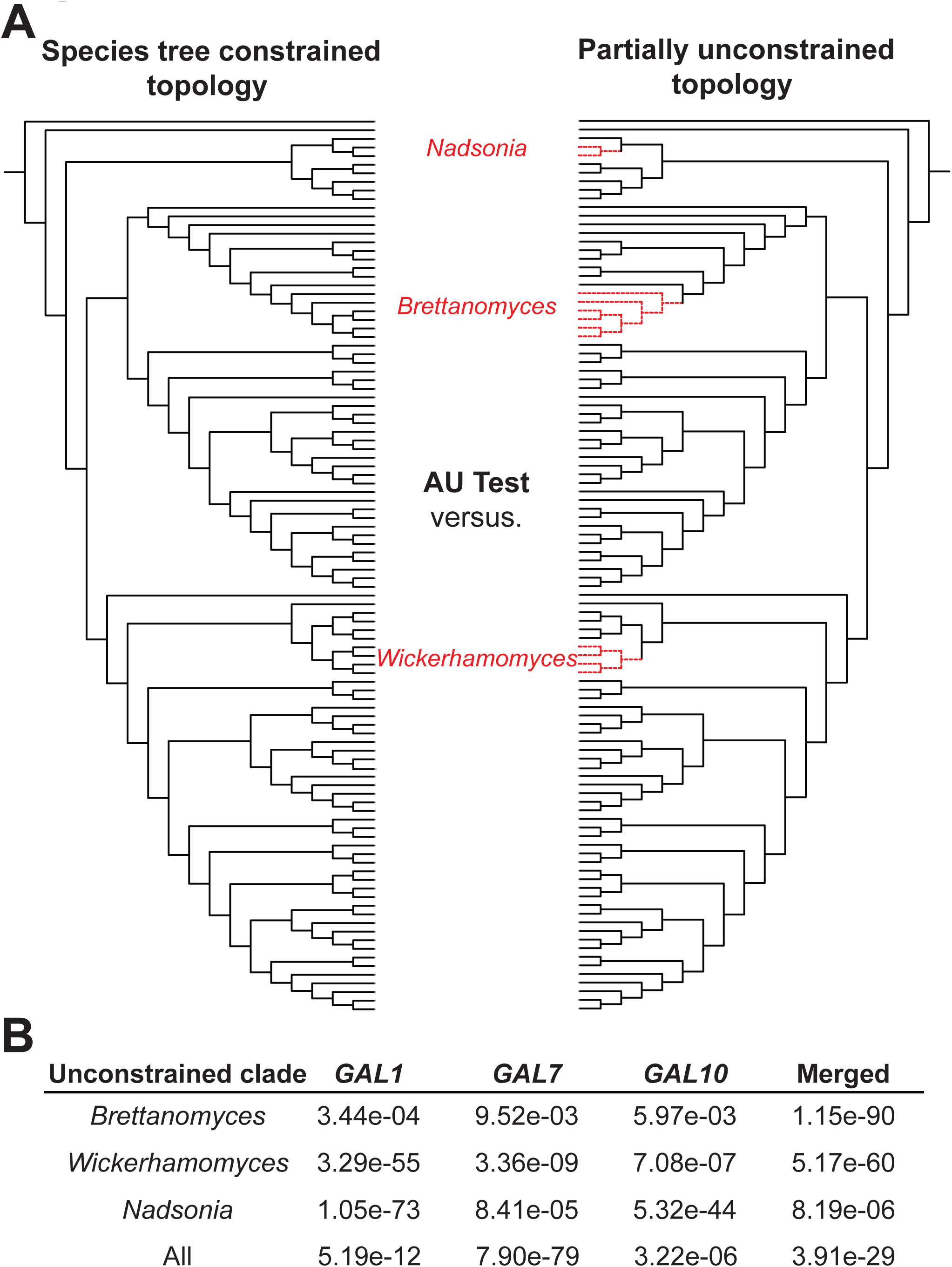
The *GAL* clusters of three lineages were acquired by HGT. (**A**) Diagrammatic representation of AU tests performed by either constraining the tree in selected lineages (as indicated in red) or not. (**B**) p-values of the AU tests are shown. All tests significantly reject their null hypotheses, indicating that the unconstrained topologies better explain the observed distribution of *GAL* genes, which is consistent with HGT as the mechanism of reacquisition.

Unexpectedly, we observed distinct *GAL* clusters in *Lipomyces starkeyi* and *Nadsonia fulvescens* (Figure 2 and Supplemental Figure 3), two species that diverged from the rest of the Saccharomycotina prior to the formation of the canonical *GAL* cluster. *L. starkeyi*, a species belonging to a lineage that is sister to the rest of the budding yeasts, contains a large gene cluster consisting of two copies of *GAL1*, a single copy of *GAL7, GALE* (predicted to only encode the epimerase domain, instead of the fused *GAL10* gene, which additionally encodes the mutarotase domain), and a gene encoding a zinc-finger domain (Supplemental Figure 3). The novel content and configuration of this cluster suggests that the *L. starkeyi GAL* gene cluster formed independently of the canonical budding yeast *GAL* cluster.

Remarkably, the structure of the *GAL* cluster of *N. fulvescens* is nearly identical to that of the CUG-Ser1 species *Cephaloascus albidus* (Figure 2 and Supplemental Figures 3 and 4), despite the fact that these two lineages are separated by hundreds of millions of years of evolution^4^. This synteny suggests that the *GAL* cluster of *N. fulvescens* was either horizontally acquired or that it independently evolved the bifunctional *GAL10* gene (fusion of galactose mutarotase (*GALM*) and UDP-galactose 4-epimerase (*GALE*) domains) and a *GAL* cluster with the same gene arrangement. Interestingly, *N. fulvescens* var. *elongata* has a pseudogenized *GAL10* gene (indicated by multiple inactivating mutations along the gene; Supplemental Figure 5), while *N. fulvescens* var. *fulvescens* has an intact *GAL10* gene, and the varieties’ phenotypes were consistent with their inferred *GAL10* functionality (Supplemental Figure 1 and Supplemental Table 3). Both varieties also contain a linked *GALE* gene, which resides ∼20 kb downstream of *GAL7*, suggesting the ongoing replacement of an ancestral *GALE*-containing *GAL* cluster by a CUG-Ser1-like *GAL* cluster containing *GAL10*. Notably, *GALE* or *GAL10* genes are present in some budding yeast species that do not utilize galactose^34^, and *N. fulvescens* var. *fulvescens* has only CUG-Ser1-like copies of the *GAL7* and *GAL1* genes required for galactose utilization. While parsimony suggests that the last common ancestor of *N. fulvescens* and its relative *Yarrowia lipolytica* was able to utilize galactose, *N. fulvescens* rests on an unusually long branch with no other known closely related species. Thus, in this case, we cannot infer whether partial cluster loss and trait loss (i.e. to the state of possessing only *GALE* and not utilizing galactose) preceded acquisition of the new functional cluster.

**Figure 5.**
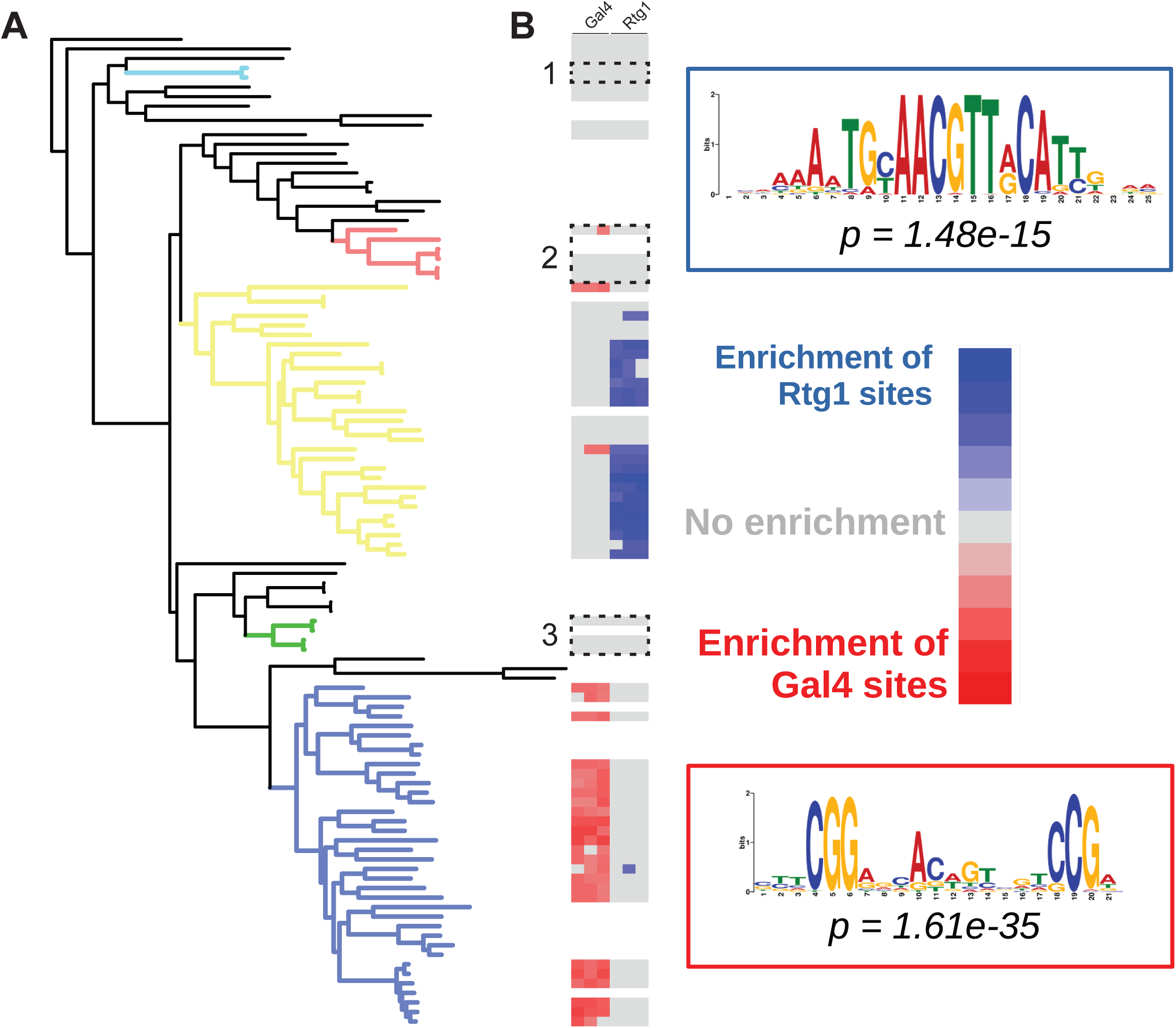
Enrichment of transcription factor-binding sites in the promoters of *GAL* enzymatic genes. (**A**) Maximum likelihood phylogeny of Saccharomycotina. Colors indicate highlighted clades: light blue – *Nadsonia*, red – *Brettanomyces*, yellow – CUG-Ser1 clade, green – *Wickerhamomyces*, and blue – Saccharomycetaceae. (**B**) Heatmap of enrichment for either Rtg1-or Gal4-binding sites in the promoters of the *GAL* genes (*GAL1, GAL10, GAL7*). White-shaded boxes indicate lineages lacking the *GAL* gene cluster.

### Allowing reacquisition is more parsimonious than enforcing loss

These synteny observations suggest three independent reacquisitions of the *GAL* cluster and at least two independent reacquisitions of the galactose utilization trait. To test the hypothesis of trait reversal, we next investigated whether, in some cases, reacquisition of the *GAL* cluster offered a more parsimonious explanation than reductive evolution. To reconcile the observed topologies of the gene and species phylogenies, we reconstructed the evolutionary events using a parsimony framework, either assuming Dollo’s law of irreversibility to be true (only gene loss was possible) or false (both gene loss and reacquisition were possible). When there was variation segregating below the species level (e.g. *N. fulvescens* and *S. kudriavzevii*^46^), we treated the species as positive for galactose utilization. When Dollo’s law was enforced, we inferred 15 distinct loss events for galactose metabolism (Figure 3A). When we allowed for the violation of Dollo’s law, we replaced a portion of the loss events with two reacquisition events, arriving at a more parsimonious inference of 11 distinct events: 9 losses and 2 reacquisitions (Figure 3A). The most parsimonious scenario did not infer trait loss for *Nadsonia*, but even adding one loss and one gain of galactose metabolism, instead of the cluster replacement scenario, still yielded a more parsimonious solution of 13 distinct events.

### Yeast *GAL* gene clusters have been horizontally transferred multiple times

From these synteny and trait reconstructions, we hypothesized that the *GAL* clusters of *Brettanomyces, Wickerhamomyces, and Nadsonia* were horizontally transferred from the CUG-Ser1 clade. This hypothesis predicts that the coding sequences of their *GAL* genes should be more similar to species in the CUG-Ser1 clade than to their closest relative possessing *GAL* genes. Thus, we calculated the percent identities of Gal1, Gal7, and Gal10 proteins between four groups of species; (A) between species in the candidate HGT recipient clade, (B) between the candidate HGT recipient clade and their closest relative with *GAL* genes, (C) between the candidate HGT recipient clade and the candidate donor clade, and (D) between the candidate HGT recipient clade and an outgroup lieage (Figure 3B, C). If the genes were vertically acquired, one would expect the percent identities to be highest in group A and then decrease in the order of group B to C to D. If the genes were acquired horizontally, then the percent identities would be higher in group C than in group B. Indeed, we found that the percent identities of the Gal proteins of group C were significantly greater than group B (Figure 3C, *p* -value = 1.79e-4). These results suggest that the *GAL* clusters of *Brettanomyces, Wickerhamomyces, and Nadsonia* were acquired horizontally from the CUG-Ser1 clade.

To further explore whether HGT occurred in these taxa, we reconstructed maximum-likelihood (ML) phylogenies for each of the *GAL* genes, as well as for the concatenation of all three (Supplemental Figures 6-9). Interestingly, we observed a consistent pattern of phylogenetic placement of *Brettanomyces, Wickerhamomyces*, and *Nadsonia GAL* genes, which grouped to different lineages than would be expected based on their species taxonomy or phylogeny (Supplemental Figure 1). The *Wickerhamomyces GAL* genes clustered with *Hyphopichia*; the *Brettanomyces GAL* genes clustered with several genera from the families Debaryomycetaceae and Metschnikowiaceae; and the *Nadsonia GAL* genes clustered with those from the family Cephaloascaceae. These observations are consistent with three independent horizontal gene transfers of *GAL* clusters into these lineages from the CUG-Ser1 clade.

**Figure 6.**
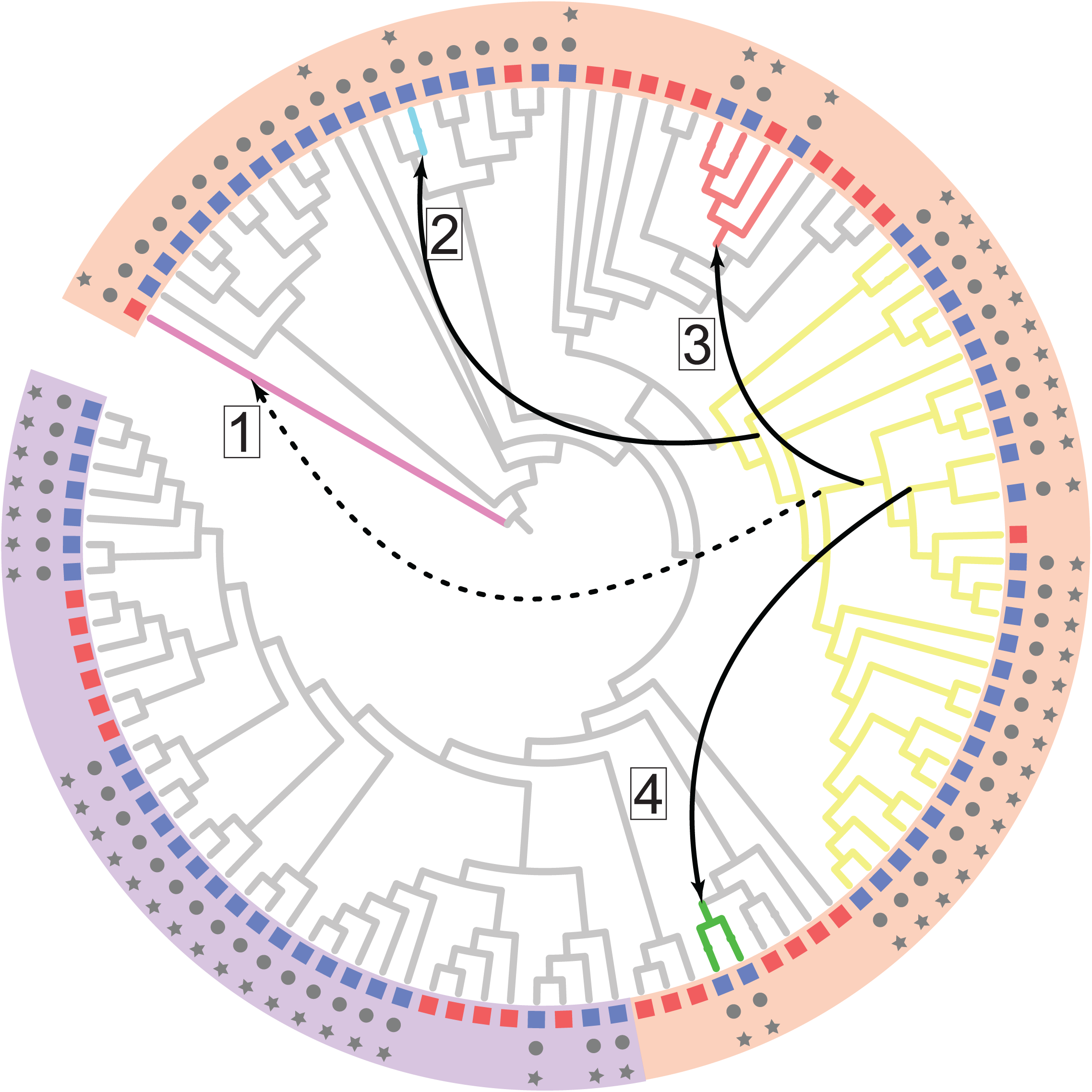
The CUG-Ser1 clade serves as a common donor of the *GAL* gene cluster to other yeasts. Cladogram of the ML phylogeny is presented with the leaf labels removed for simplicity. The colored boxes represent the species’ ability to utilize galactose (blue = positive/variable, red = negative), gray circles indicate the presence of a full set of *GAL* enzymatic genes, and gray stars indicate that those *GAL* genes are clustered. Five lineages on the cladogram are colored: pink - *Schizosaccharomyces pombe* (a member of the subphylum Taphrinomycotina with a transferred *GAL* cluster that does not confer utilization), light blue – *Nadsonia*, red – *Brettanomyces*, yellow – CUG-Ser1 clade, and green – *Wickerhamomyces*. Numbered boxes and arrows depict the four horizontal transfer events of the *GAL* cluster. The colored arcs encompassing the cladogram represent the predicted regulatory mode of the *GAL* genes: orange – Rtg1/Rtg3 (non-Gal4) and purple – Gal4.

To formally test the hypothesis of *GAL* HGT, we used Approximately Unbiased (AU) tests (Figure 4A). Specifically, we generated multiple maximum likelihood phylogenetic trees using alignments of *GAL* genes with constraints on the placements of various taxa: (i) fully constrained to follow the species tree, (ii) unconstrained in the *Brettanomyces* lineage, (iii) unconstrained in the *Wickerhamomyces* lineage, (iv) unconstrained in the *Nadsonia* lineage, and (v) unconstrained in all three candidate HGT lineages (*Brettanomyces, Wickerhamomyces*, and *Nadsonia*). By comparing the partially constrained trees to the fully constrained tree with AU tests, we found that each of the proposed horizontal transfer events was statistically supported (Figure 4B). These results were consistent across individual alignments of the *GAL* genes and when all three lineages were examined together (Figure 4B). From these results, we conclude that the *GAL* clusters of the *Brettanomyces, Wickerhamomyces*, and *Nadsonia* lineages were likely acquired via HGT from ancient CUG-Ser1 yeasts.

### Regulatory mode correlates with the horizontal gene transfers

Gal4 is the key transcriptional activator of the *GAL* cluster in *S. cerevisiae* and responds to galactose through the co-activator Gal3 and co-repressor Gal80. This mode of regulation is thought to be restricted to the family Saccharomycetaceae and is absent in other yeasts and fungi^47^. In other budding yeasts (including *C. albicans*, the most thoroughly studied CUG-Ser1 species, as well as *Y. lipolytica*, an outgroup to *S. cerevisiae* and *C. albicans*), regulation of the *GAL* cluster is thought to be under the control of the activators Rtg1 and Rtg3^48^. These two regulatory mechanisms respond to different signals and have dramatically different dynamic ranges. In Gal4-regulated species, the *GAL* cluster is nearly transcriptionally silent in the presence of glucose and is rapidly induced to high transcriptional activity when only galactose is present. In contrast, Rtg1/Rtg3-regulated species have high basal levels of transcription and are weakly induced in the presence of galactose^48^.

Intriguingly, all putative donor lineages of the *GAL* genes were from the CUG-Ser1 clade of yeasts, and no transfers occurred from or into the family Saccharomycetaceae. To examine whether the relaxed Rtg1/Rtg3 regulatory regimen of the CUG-Ser1 yeasts might have facilitated their role as an HGT donor, as opposed to the Gal4-mediated regulation of the Saccharomycetaceae, we identified sequence motifs that were enriched 800 bp upstream from the coding regions of the *GAL1, GAL7*, and *GAL10* genes (Supplemental Table 4). Then, based on the existing experimental evidence on the regulation of the *GAL* genes^23,24,48^, we divided the yeast species into Saccharomycetaceae and non-Saccharomycetaceae species. We then ran a selective motif enrichment analysis to determine if any regulatory motifs were enriched in one group, but not the other. We found that the top enriched motifs corresponded to the known Gal4-binding site in the Saccharomycetaceae^20^ and the known Rtg1-binding site in the non-Saccharomycetaceae species^48^ (Figure 5A and B, Supplemental Table 4), consistent with the previously documented regulatory rewiring of the *GAL* genes that occurred at the base of the family Saccharomycetaceae^48^. In general, the enrichment of Rgt1-binding sites was patchier and did not include the HGT recipient lineages, the previously characterized Rtg1-regulated *GAL* cluster of *Y. lipolytica*^48^, or several CUG-Ser1 clade species (e.g. *Ce. albidus*).

Taken together, our new results suggest that the switch to the Gal4-mode of regulation, which is tighter and involves multiple unlinked and dedicated regulatory genes, reduced the likelihood of horizontal transfer into naïve genomes or genomes that had lost their *GAL* pathways. Specifically, any *GAL* cluster regulated by Gal4 would not be able to be transcribed or properly regulated if it were horizontally transferred into a species lacking *GAL4* and other regulatory genes. In contrast, Rtg1 and Rtg3 are more broadly conserved, and any horizontally transferred *GAL* cluster regulated by them would likely be sufficiently transcriptionally active, providing an initial benefit to the organism.

## Discussion

Budding yeasts have diversified from their metabolically complex most recent common ancestor over the last 400 million years^2,4^. While they have evolved specialized metabolic capabilities, their evolutionary trajectories have been prominently shaped by reductive evolution^2,4,49,50^. Here, we present evidence that losses of the *GAL* genes and galactose metabolism in some lineages were offset, tens of millions of years after their initial losses, by eukaryote-to-eukaryote horizontal gene transfer (Figure 6). While reacquired ancestral traits have been documented in several eukaryotic lineages, our observation of galactose metabolism reacquisition differs in a few regards. First, the majority of reported events did not identify the molecular mechanism or the genes involved in the reacquired traits. Second, few studies have comprehensively sampled taxa and constructed robust genome-scale phylogenies onto which the examined traits were mapped, a requirement for robustly inferring trait evolution. Remarkably, we observed trait reversal in at least two independent lineages, with a third possible lineage, suggesting that the recovery of lost eukaryotic metabolic genes may be an important and underappreciated driver in trait evolution in budding yeasts, and perhaps more generally in fungi and other eukaryotes. In line with our study, budding yeasts also have reacquired lost metabolic traits from bacteria, supporting the hypothesis that regains via HGT offset reductive evolution^44^.

The dearth of HGT from Saccharomycetaceae into other major clades provides clues into the potential limits on ancestral trait reacquisition via HGT. We propose the transcriptional rewiring to Gal4-mediated regulation imposed a restriction on the potential for benefit of transferred *GAL* clusters. Since Gal4-mediated gene activation is tightly coordinated and the off-state is less leaky^47^, any transferred *GAL* cluster lacking Gal4-binding sites into a species with exclusively Gal4-mediated activation in response to galactose would not be able to activate the transferred genes. Similarly, transfer of a Gal4-regulated gene cluster into a species lacking *GAL4* and other upstream regulators would have limited potential for activation. For the case of transfer between two species whose regulation does not rely on Gal4, the transferred *GAL* cluster would be transcriptionally active because the broadly conserved transcription factors Rtg1 and Rtg3 could further enhance moderate basal transcriptional activity^48^. Thus, even leaky levels of transcription would provide a benefit in the presence of galactose that could further be refined, possibly to become regulated by lineage-specific networks. Under this model, the likelihood of HGT is partly determined by the potential activity of the transferred genes and by the recipient’s ancestral regulatory mode.

More generally, our findings demonstrate that reductive evolution is not always a dead end, and gene loss can be circumvented by HGT from distantly related taxa. However, the scope of genes that can be regained in this fashion is likely limited. In particular, the *GAL* genes of the CUG-Ser1 clade of budding yeasts represent something of a best-case scenario. First, all enzymatic genes needed for phenotypic output are encoded in a cluster, facilitating the likelihood that all necessary genes for function are transferred together^32,51^. Second, the regulatory mode of these *GAL* genes is conducive to function in the recipient species, as they are loosely regulated by conserved factors with moderate basal activity. Third, the genes would provide a clear competitive advantage in environments with galactose.

The modern interpretation of Dollo’s law is that species cannot return to a previous character state after loss. Alongside recently reported character state reversals in petunias after pseudogene reactivation^52^, our results of reacquisition of galactose metabolism and *GAL* genes by HGT can be considered a case of character state reversal. However, the previous example fits into the model that, for groups undergoing adaptive radiations, lost traits seem to “flicker” on and off, resulting in an unusual distribution of character states on the phylogeny. Here, and in the recently described reacquisition of alcoholic fermentation genes from bacteria in fructophilic yeasts^44^, the ancestral genes were completely lost from the genome, and they were restored far later than could be explained by the flickering of traits during adaptive radiations. The reacquisition of galactose metabolism in budding yeasts represents a striking example of gene and trait reversal by eukaryote-to-eukaryote horizontal gene transfer and provides insight into the mechanisms by which Dollo’s law can be broken.

## Methods

### *GAL* gene identification

We analyzed 96 publicly available genome sequences used in a recent study of the Saccharomycotina phylogeny^3^ (86 Saccharomycotina, 10 outgroups), as well ten additional species belonging to clades where we identified potentially deep losses of the *GAL* gene cluster. Of the latter ten species, five genome sequences, including *Nadsonia fulvescens* var. *fulvescens*, were published recently^4^, while genome sequences for five new species are published here. Due to their importance to this study and since previously published genome sequences may have been from different strains that were unavailable for phenotyping, eight additional genome sequences were generated for taxonomic type strains. In total, 104 genome sequences were analyzed. All genome sequences generated after a backbone phylogeny was compiled from data published before 2016^3^ are denoted Y1000+ in Supplemental Figures 6-9. The presence of *GAL* genes in the genome assemblies was inferred with TBLASTN^53^ v2.7.1 using the *C. albicans* Gal1, Gal7, and Gal10 sequences as queries, followed by extraction of the open reading frame centered on the location of the best hit. The structure and synteny of the clusters were manually curated and documented. For *S. kudriavzevii*, where balanced variation is segregating for the *GAL* pathway^46^, phylogenetic analyses were performed with the taxonomic type strain (cannot grow on galactose), whereas summary figures (Supplemental Figure 3 and Figures 1, 3, and 6) show a reference strain (ZP591) that can grow on galactose.

### Sequencing and assembly of genomes

For the five new genomes sequenced here, genomic DNA was sonicated and ligated to Illumina sequencing adaptors as previously described^26^. The paired-end library was sequenced on an Illumina HiSeq 2500 instrument, conducting a rapid 2×250 run. To generate whole genome assemblies, paired-end Illumina reads were used as input to a meta-assembler pipeline iWGS^54^. The quality of the assemblies was assessed using QUAST^55^ v3.1, and the best assembly for the newly described species was chosen based on N50 statistics.

### *GAL* gene similarity analysis

To calculate the percent identities between Gal proteins, we first aligned the protein sequences for each species (see Supplemental Table 1 for species used) of Gal1, Gal7, and Gal10 and generated percent identity matrices using Clustal Omega^56^. These results were then subdivided into four groups: (1) the percent identities between species within the potential HGT recipient clade, (2) the percent identities between species of the recipient clade and their closest relative with *GAL* genes, (3) the percent identities between species of the recipient clade and species in the donor lineage, and (4) the percent identities between species of the recipient clade and an outgroup lineage (i.e. *S. cerevisiae*). Next, a similarity score was calculated by normalizing the percent identity values of each group to the average value of the fourth group:

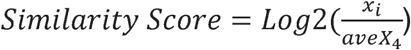

### Phylogenetic analyses

Sequence alignments were conducted using MAFFT^57^ v 7.409 run in the “--auto” mode. Alignments were subjected to maximum-likelihood phylogenetic reconstruction using RAxML^58^ v8.1.0 with 100 rapid bootstrap replicates. Constrained phylogenetic trees were generated with RAxML using the “-g” option, with the constraint tree identical to the species tree, except for the species/lineage of interest, whose position on the tree was allowed to be optimized by the ML algorithm. Statistical support for the HGT events involving *GAL* genes was determined using the Approximately Unbiased (AU) test, by comparing the various partially constrained ML phylogenies and the fully constrained phylogeny. The AU test was performed with IQ-TREE^59^ v1.6.8 (-au option), which was run with the General Time Reversible model, substitution rate heterogeneity approximated with the gamma distribution (-m GTR+G), and with 10,000 replicates (-zb 10000)

### Regulatory motif enrichment

Sequences of 800 bp upstream of the start codon of all identified *GAL* genes were extracted and subjected to a regulatory motif identification analysis using MEME^60^ v5.0.2, with the following constraints: maximum number of motifs = 20 (-nmotifs 20), maximum length of motif = 25 bases (-maxw 25), any number of motif repetitions (-anr), active search of reverse complement of the used sequence (-revcomp), and the log-likelihood ratio method (-use_llr). Selective enrichment of motifs was determined by splitting the sequences into Saccharomycetaceae and non-Saccharomycetaceae groups and running AME^61^ v5.0.2, with each group being the control group in one analysis and the test group in a second analysis.

### Species tree reconstruction

Our data matrix was composed of 104 budding yeasts and 10 outgroups, comprising of 1,219 BUSCO genes (601,996 amino acid sites); each gene had a minimum sequence occupancy ≥57 taxa and sequence length ≥167 amino acid residues. For the concatenation-base analysis, we used RAxML version 8.2.3 and IQ-TREE^59^ version 1.5.1 to perform maximum likelihood (ML) estimations under an unpartitioned scheme (a LG+GAMMA model) and a gene-based partition scheme (1,219 partitions; each gene has its own model), respectively. As a result, four ML trees produced by two different phylogenetic programs and two different partition strategies were topologically identical. Branch support for each internode was evaluated with 100 rapid bootstrap replicates using RAxML^62^. For the coalescence-based analysis, we first estimated individual gene trees with their best-fitting amino acid models, which were determined by IQ-TREE^59^ (the “–m TESTONLY” option); we then used those individual gene trees to infer the species tree implemented in the ASTRAL program^63^, v4.10.2.

The reliability for each internode was evaluated using the local posterior probability measure^64^. Finally, internode certainty (IC) was used to quantify the incongruence by considering the most prevalent conflicting bipartitions for each individual internode among individual gene trees^65,66,67^ implemented in RAxML^58^ v8.2.3. The relative divergence times were estimated using the RelTime^68^ in MEGA7^69^. The ML topology was used as the input tree.

### Growth assays

We previously published galactose growth data for the majority of species^4,70^.Growth experiments were performed for an additional nine species separately (Supplemental Table 3). All species were struck onto yeast extract peptone dextrose (YPD) plates from freezer stocks and grown for single colonies. Single colonies were struck onto three types of plates minimal media base (5g/L ammonium sulfate, 1.71g/L Yeast Nitrogen Base (w/o amino acids, ammonium sulfate, or carbon), 20g/L agar) treatments with either: 2% galactose, 1% galactose, or 2% glucose (to test for auxotrophies). We also re-struck the specific colony onto YPD plates as a positive control. All growth experiments were performed at room temperature. After initial growth on treatment plates, growth was recorded for the first round, and we struck colonies from each treatment plate onto a second round of the respected treatment to ensure there was no nutrient carryover from the YPD plate. For example, a single colony from 2% galactose minimal media plate was struck for a second round of growth on a 2% galactose minimal media plate. We inspected plates every three days for growth for up to a month. Yeasts were recorded as having no growth on galactose if they did not grow on either the first or second round of growth on galactose.

## Acknowledgments

We are grateful to Carlos A. Rosa for providing the strain *Wickerhamomyces* sp. UFMG-CM-Y6624. We thank the Rokas and Hittinger labs for comments and discussions and the University of Wisconsin Biotechnology Center DNA Sequencing Facility for providing Illumina sequencing facilities and services. This material is based upon work supported by the National Science Foundation under Grant Nos. DEB-1442113 (to A.R.) and DEB-1442148 (to C.T.H. and C.P.K.), in part by the DOE Great Lakes Bioenergy Research Center (DOE BER Office of Science DE-SC0018409 and DE-FC02-07ER64494 to Timothy J. Donohue), USDA National Institute of Food and Agriculture (Hatch Project 1020204), and National Institutes of Health (NIAID AI105619 to A.R.), and a Guggenheim fellowship (to A.R). C.T.H. is a Pew Scholar in the Biomedical Sciences, a Vilas Early Career Investigator, and a H. I. Romnes Faculty Fellow, supported by the Pew Charitable Trusts, Vilas Trust Estate, and Office of the Vice Chancellor for Research and Graduate Education with funding from the Wisconsin Alumni Research Foundation, respectively.

## Data deposition

Raw DNA sequencing data were deposited in GenBank under Bioproject ID PRJNA647756. Whole Genome Shotgun assemblies have been deposited at DDBJ/ENA/GenBank under the accessions XXX-XXXX (pending processing). The versions described in this paper are versions XXX-XXXX (pending processing).

## Author contributions

M.A.B.H. (study design, preliminary phylogenetic analyses, sequence analyses, cluster analyses, text); J. K. (study design, genome assemblies, phylogenetic analyses, motif enrichment analyses, text); D.A.O. (genomic DNA isolation, library preparation, yeast growth assays); X.S. (phylogenomic analyses); A.L.L. (cluster analyses); X.Z. (preliminary genome annotations and analyses); J.DeV., A.B.H. (genomic DNA isolation, library preparation); C.P.K. (support and supervision, study design); and A.R., and C.T.H. (support and supervision, study design, text).

## Supplemental Figures and Tables

Supplemental Table 1. Strains used in this study.

**Table S2.**
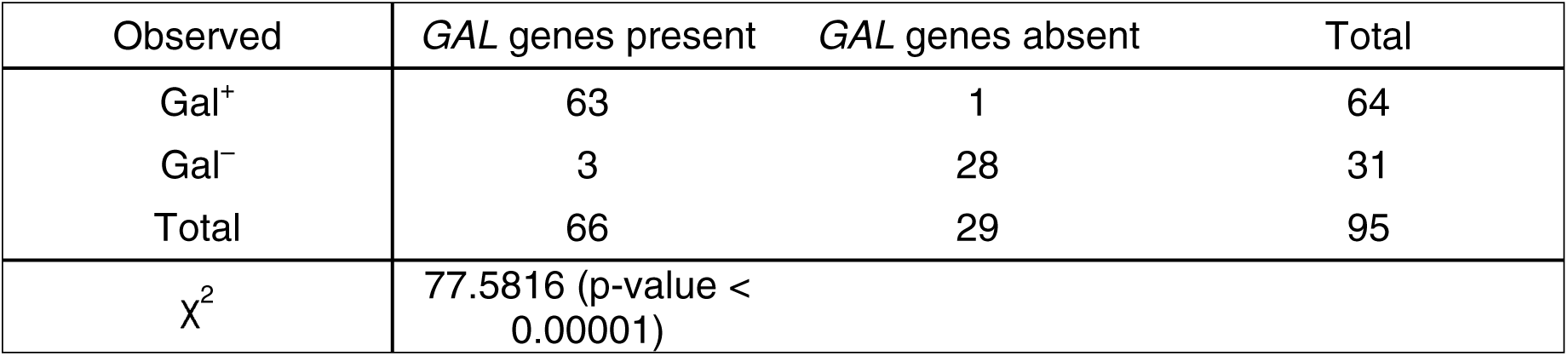
Chi-squared (χ^2^) test of genotype-to-phenotype associations of species presented in Supplemental Figure 3. We used our phenotypes in cases where our data disagreed with *The Yeasts* book^2^.

| Observed | GAL genes present | GAL genes absent | Total |
| --- | --- | --- | --- |
| Gal<sup>+</sup> | 63 | 1 | 64 |
| Gal<sup>-</sup> | 3 | 28 | 31 |
| Total | 66 | 29 | 95 |
| $\chi^2$ | 77.5816 (p-value < 0.00001) | | |

**Table S3.**
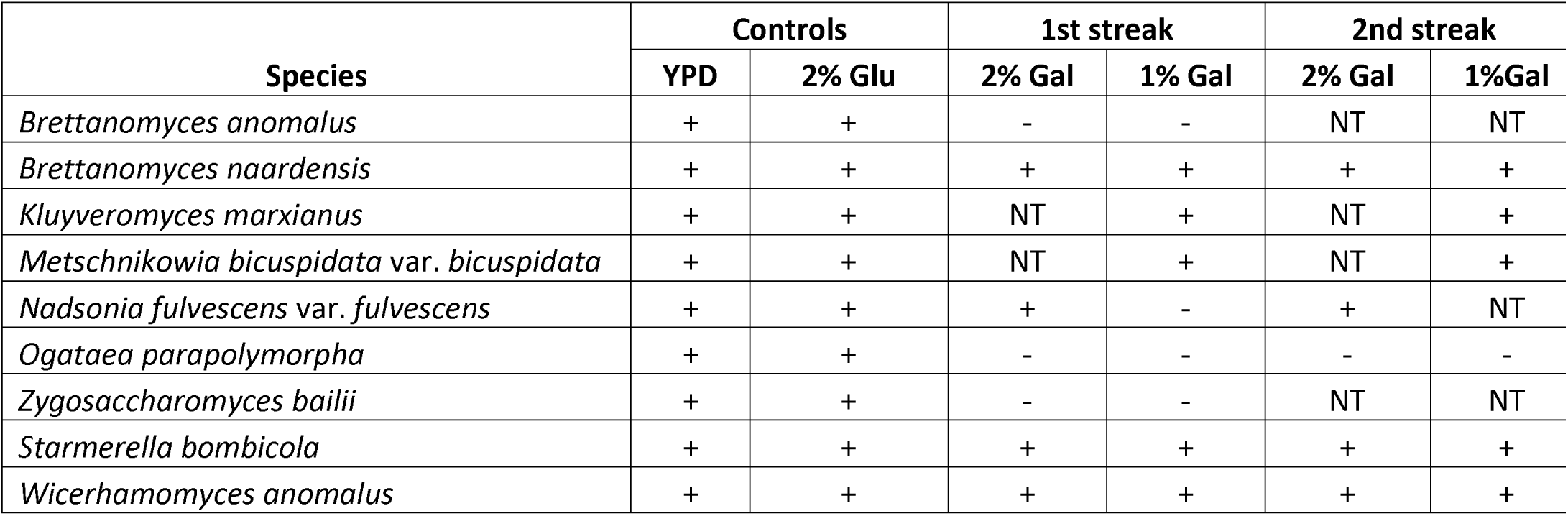
Galactose growth phenotyping of key species. NT, not tested.

Supplemental Table 4. Per-species p-values for the presence of Gal4-and Rtg1-binding site motifs in individual *GAL* genes.

Supplemental Figure 1. Genome-scale maximum likelihood phylogeny.

Supplemental Figure 2. Genome-scale internode certainty cladogram.

Supplemental Figure 3. Distribution of the structure of *GAL* gene clusters. Both cluster structure and growth characteristics are mapped onto the relative divergence timetree. Growth on galactose is indicated by the colored squares next to each species (green=blue, yellow=variable, red=negative). Asterisks next to certain species names indicated either a new genome sequence published here (**) or an additional genome sequence from a recent study (*)^4^, including *Nadsonia fulvescens* var. *fulvescens*. To ensure phenotyping could be performed on sequenced strains, we also sequenced the genomes of the taxonomic type strains for eight species and report those *GAL* clusters here (^). The syntenic structure of the *GAL* genes are displayed to the right of the growth characteristics for each species. The structure of the *Nadsonia fulvescens* var. *elongata* cluster is shown in Supplemental Figure 4.

Supplemental Figure 4. Surprisingly syntenic *GAL* clusters between diverse lineages.

Supplemental Figure 5. Alignment of the *GAL10* genes of *N. fulvescens* var. *fulvescens* and *N. fulvescens* var. *elongota*. Genes were aligned using MAFFT v 7.409 using -- auto. Likely inactivating mutations are shown in various colors: mutation of the start codon in orange, frameshift mutations in blue, in-frame nonsense mutations in red, and insertions in green. One in-frame deletion is shown in purple.

Supplemental Figure 6. Gene tree of *GAL1* genes.

Supplemental Figure 7. Gene tree of *GAL7* genes.

Supplemental Figure 8. Gene tree of *GAL10 genes*.

Supplemental Figure 9. Concatenated gene tree of the *GAL*actose enzymatic gene cluster.

Supplemental Figure 10. Percent identities of *GAL* genes as calculated by the comparisons shown in Figure 3.

