## Supplemental Figures 1-10 for "Repeated horizontal gene transfer of *GAL*actose metabolism genes violates Dollo’s law of irreversible loss"

### Supplemental Figure 1

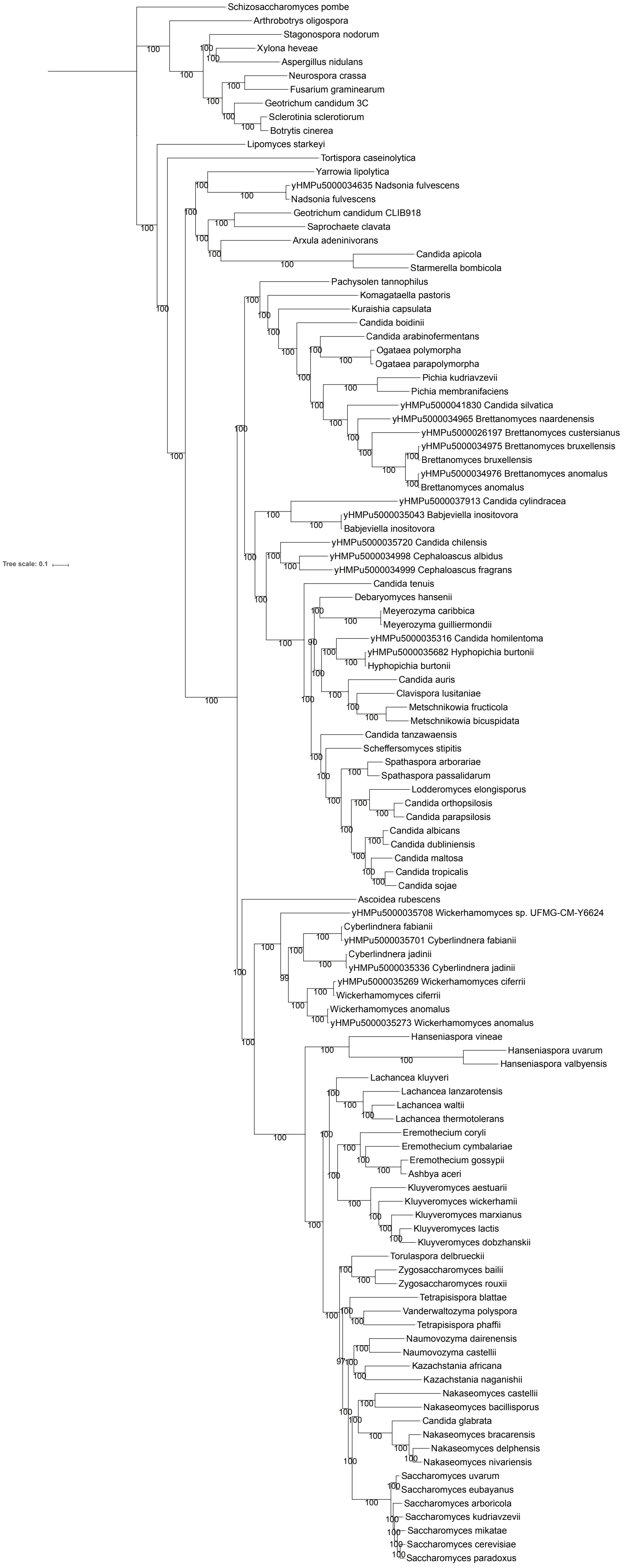

### Supplemental Figure 2

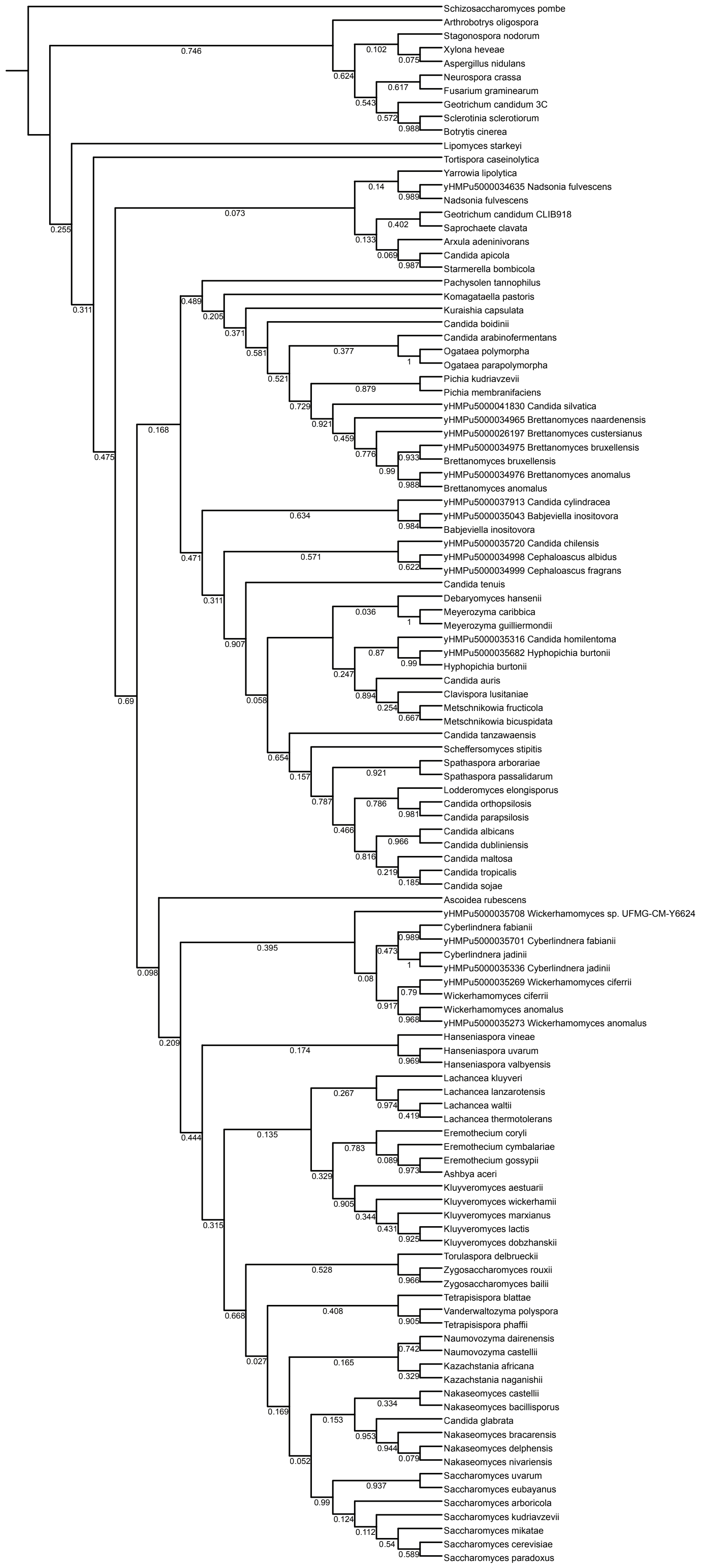

### Supplemental Figure 3

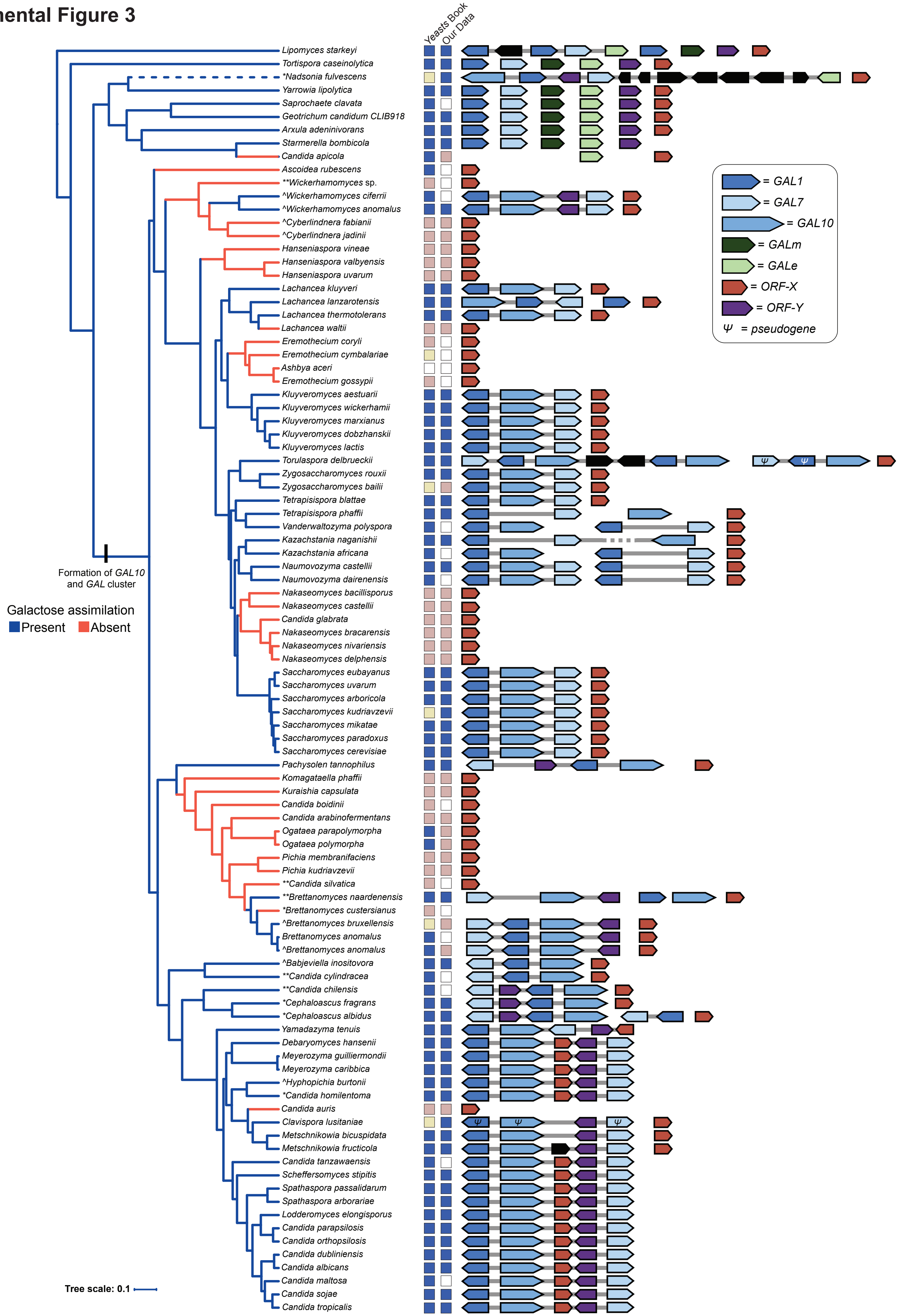

### Supplemental Figure 4

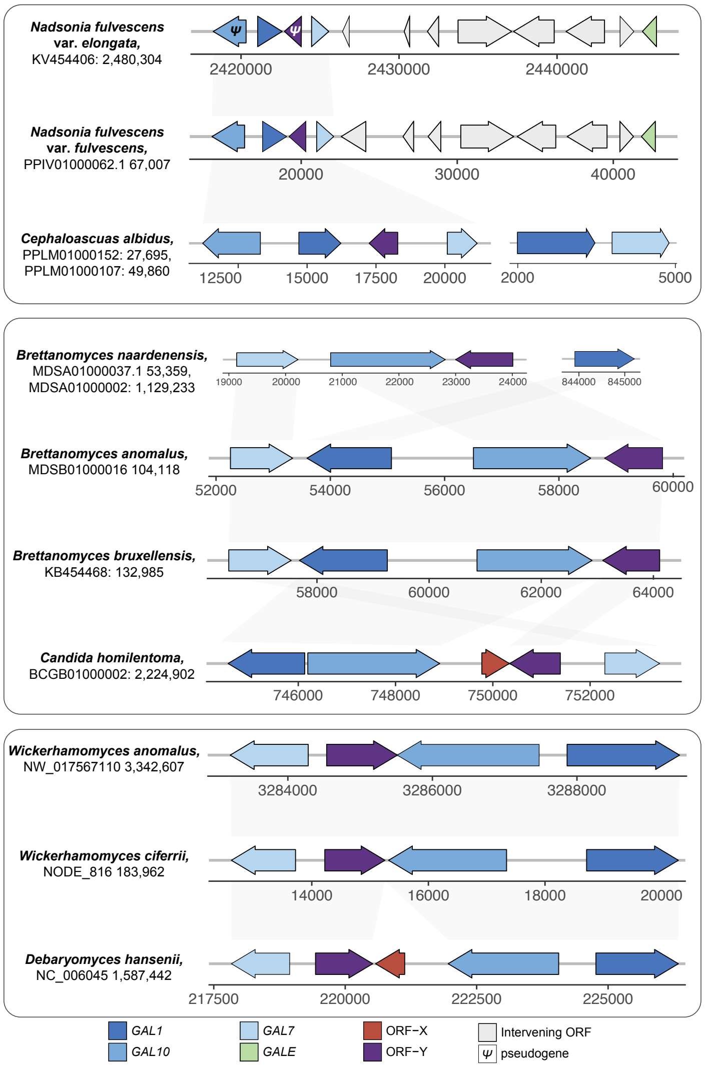

#### Supplemental Figure 5

```
fulvescens    1 ATGTCTTATGTATTAGTTACCGGCGGTGCTGGTTATATCGGCTCCCATACAGTGGTTGAG
elongota      1 ..A..C....................C..C..............................

fulvescens   61 CTTTTCAACAATGGCTACAGCGTAGTCATTGTGGATAACTTGGTCAATTCCTCCTACGAC
elongota     61 .-..........A....-.A.........C....-.........................

fulvescens  121 TCAGTAGCAAGACTAGAAGTCTTGTGTAAGCGGAAGATTCCCTTTTTCAAAATCGACACT
elongota    118 .....G..C.................................-..GAAG.......T...

fulvescens  181 AGTGACGCCATAGCTCTGGATGCCGTCTTCAAGCAGTACAAGATCGACTCAGTCTTGCAC
elongota    177 .....................A..........A..A.........A..............

fulvescens  241 TTTGCCGCTTTAAAGGCCGTTGGCGAATCGACCAAGATTCCTCTTGAATACTATGAAAAC
elongota    237 .................TA.AA.T.....A...........C..................

fulvescens  301 AACGTTGGTGGCACTATTGCTTTATTAAAAGTGATGCGACAAAACAATTGTAAAAAAATT
elongota    297 ..T...A.C.....C.................A......T...........T.......

fulvescens  361 GTGTTTTCTTCCTCGGCTACCGTCTACGGAGATGCTACGCGATTCCCCGATATGATACCT
elongota    357 ..............A...........................................T.

fulvescens  421 ATTCCTGAACACTGTCCTACCGGCCCTACCAACCCTTATGGCATGACAAAAATTACCATC
elongota    417 .........T.......C.....T..C..G.......................C......

fulvescens  481 GAGAACATCATTAGAGATCTCTTCAGTAGCGACAAATCATGGAAGTCGGCTATTTTGAGA
elongota    477 ...................................C..C.....................

fulvescens  541 TATTTTAATCCCATTGGCGCTCACCCATCTGGGTTAATTGGCGAAGATCCTTTGGGTATC
elongota    537 .GC.A...........................CC..G....T..G.....C.....C...

fulvescens  601 CCCAACAACTTATTGCCCTTCTTAGCCCAGGTTGCCATTGGCCGCCGAGAGAAGATGTGC
elongota    597 .....T........A...........T........T..........A......A....A.

fulvescens  661 ATTTTTGGTGATAATTATGATTCTAAAGATGGAACTCCCATTCGTGATTATATACATGTA
elongota    657 ............G...G......C.G...........................C..C..G

fulvescens  721 GTTGATTTGGCCAAGGGCCATATTGCGGCTCTTGACCACTTGAACGAATCGCAGCAAGGA
elongota    717 ..............A.A......C..C..C...A.TT....A..T.....A........G

fulvescens  781 TTATGCCGTGAATGGAACTTGGGTACAGGTAACGGTTCTACCGTAATCGAAGTCTTCAAC
elongota    777 ......................................C..............T......

fulvescens  841 GCTTTCTGTAAGGCCGTAGGTAGAGACTTGCCGTTTGAAATCGCCGGTCGCCGAGCCGGT
elongota    837 ...............A.....C....TC.A..C........TA....C.......AT...

fulvescens  901 GATGTACTAAACCTTACGGCAGACGCTAGAAGAGCGAACAAAGAGTTGAAATGGCATGCG
elongota    897 A.C.............................G..T......A............TCA.A

fulvescens  961 GAATTATCCATTGATGATGCCTGTAAGGATTTGTGGAAATGGACAACTGATAATCCTATG
elongota    957 ......C...C.......A.....GGA.................CG..............

fulvescens 1021 GGGTTTCAAATTGAAGATTACAAATGGAAGCACTTCAATAATGAATCCCATTTTGAAGAT
elongota   1017 .....C.........A.-...G......G.T.............................

fulvescens 1081 AGATTACATACTTTAATTTCCAGTGACGGAAAGTTTCAATTCTCCGTTAGCAATCTGGGA
elongota   1076 ......T.......G...........T..T.G....G...................T...

fulvescens 1141 GCTAGTGTTGTGGATGCGTCTCTGGACGGTGTTAAACTGTGCTGTGGTTTCGATAATGAA
elongota   1136 .........................GT........G..A....A.A....T.........

fulvescens 1201 GAAGGTTATTTGAGAAAGGATAACCCTTTCTTTGGCGCCACCATTGGCAGGGTTGCCAAT
elongota   1196 ........C..A....................CA........G.......AAA..-A...

fulvescens 1261 AGAGTCAAGGGTGGTAAATTAACTGTTAACGGCTCTACTTATCAGCTACCCCTAAATGAA
elongota   1255 ..............C...........C..A...............T.............G

fulvescens 1321 AATGGTGTCAATACTATTCATGGTGGGTTATCCGGATTCGACAAAAAACTTTTCTTGGGT
elongota   1315 ...A........................C.....A...........G............G

fulvescens 1381 CCAATTGTTCGTAAGAATCTCCAAACTGATTTAAACCAAATGGAATTTTTACTTGTTGAC
elongota   1375 T.C................CT.......GC...........AT..........A.....

fulvescens 1441 AAAGATGGTGAAAACAACTTCCCAGGTGATTTAGAGGTCAAGGTAATCTACACTCTTCAA
elongota   1435 G.G..G........T..T...........C....TC...........T...T........

fulvescens 1501 AAGGTCGAAAAGGGTGGCGCCATCGGCTTGGAATACGAAGCTAAATTATTGCCTGATACT
elongota   1495 ..A..T...........T..T.A......A...................C....A.....

fulvescens 1561 AATGTCCATGAAACTGCCGTTTCCATTACCAATCATACCTATTGGAACATCGGGAATAAC
elongota   1555 ...A...CC.............T.............T.T........T............

fulvescens 1621 TCTACTATAGAGGGTACTGAAATCGCTTTAATTACCGATAAACATTTGCAGGTTGGAGCT
elongota   1615 ............A.C..---GG....C..T............A.......A...ATG...

fulvescens 1681 GATTTATTGCCGACCGGGGATATAATCATCAATAATAATATCGCCAGGATTGAATCCGGT
elongota   1672 ......GC.TT......----C.T..............G.......CT......A.....

fulvescens 1741 AAATTTACTACTTTGAATCAAAAATCTCCTGAATTTGACTATTGTTTTACTGTTGCGGAT
elongota   1728 ....C.....TC...GG.............C.....T....C..C.....C...A.....

fulvescens 1801 CCC---AAGAATTTTCAAATCGATACCAGGTCCAAGAATATGCAAGTTATCGCTAAGGCT
elongota   1788 ...A--..A..A...........CC....A.......TC.....................

fulvescens 1860 TACCATCCTGAAACCAAGATCCGCTTTATAACCTCTACAACTGAACCGTCATTCCAATTT
elongota   1848 ...........G..........A.........T..............C......T..C..

fulvescens 1920 TACACTGGTGACCATATTAATGTTGAAGGTTTATTTGGAAAGCGTGCTGGATTTTGCTTA
elongota   1908 ........A........C...........C..................A.C.........

fulvescens 1980 GAGGCTGCTAGGTTTATTGATTCCTGTAACCACGAAGATTGGCGGGCAGGTATGGTATTA
elongota   1968 .....C.....A.................T......C......A.........C......

fulvescens 2040 AAGAATGGGGAAGTCTACGGTTCAAAGACCCTATACAGTTTCAGCACTGAATAG
elongota   2028 .....G...................GT..T..G..G..C....A....A.....
```

#### Supplemental Figure 6

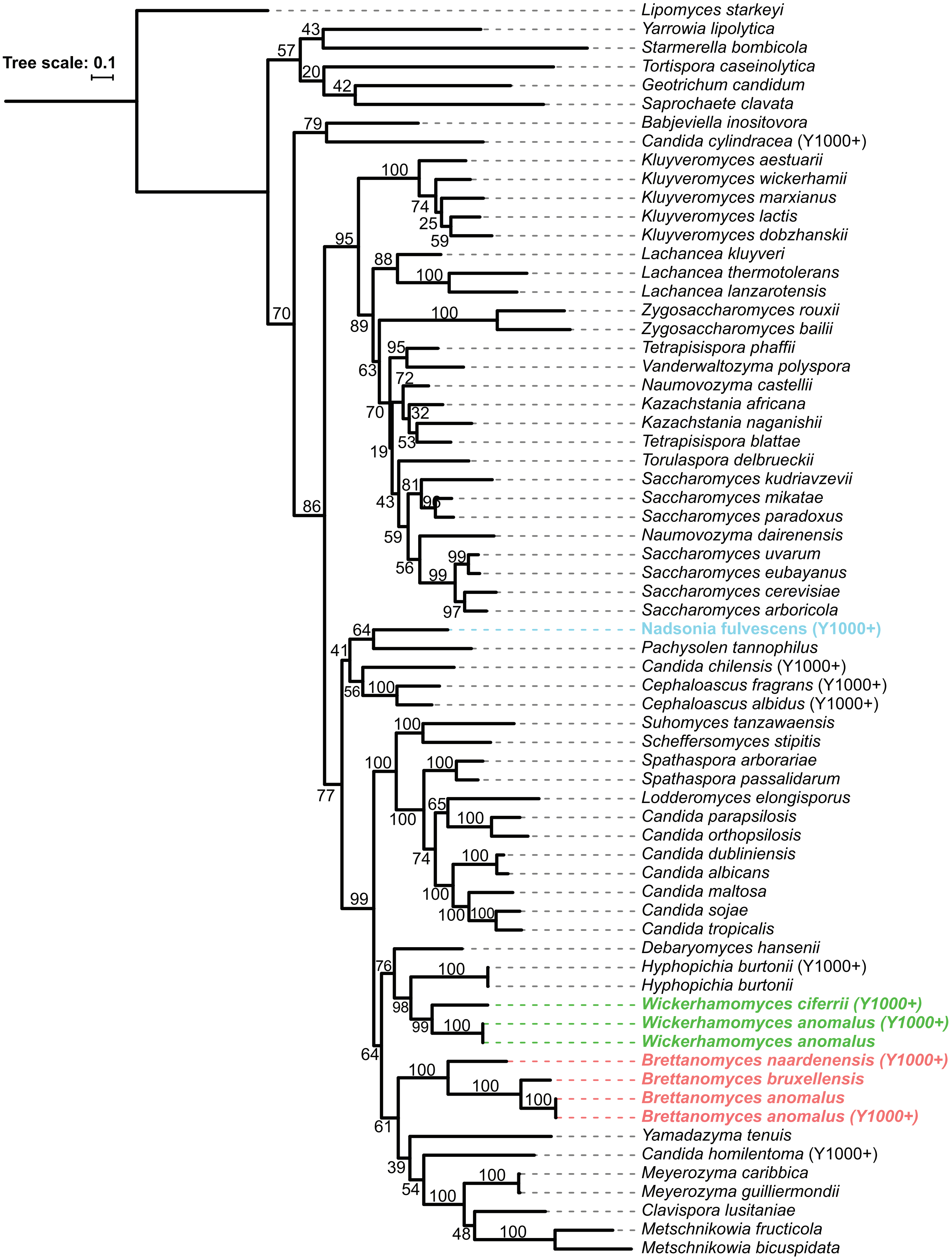

#### Supplemental Figure 7

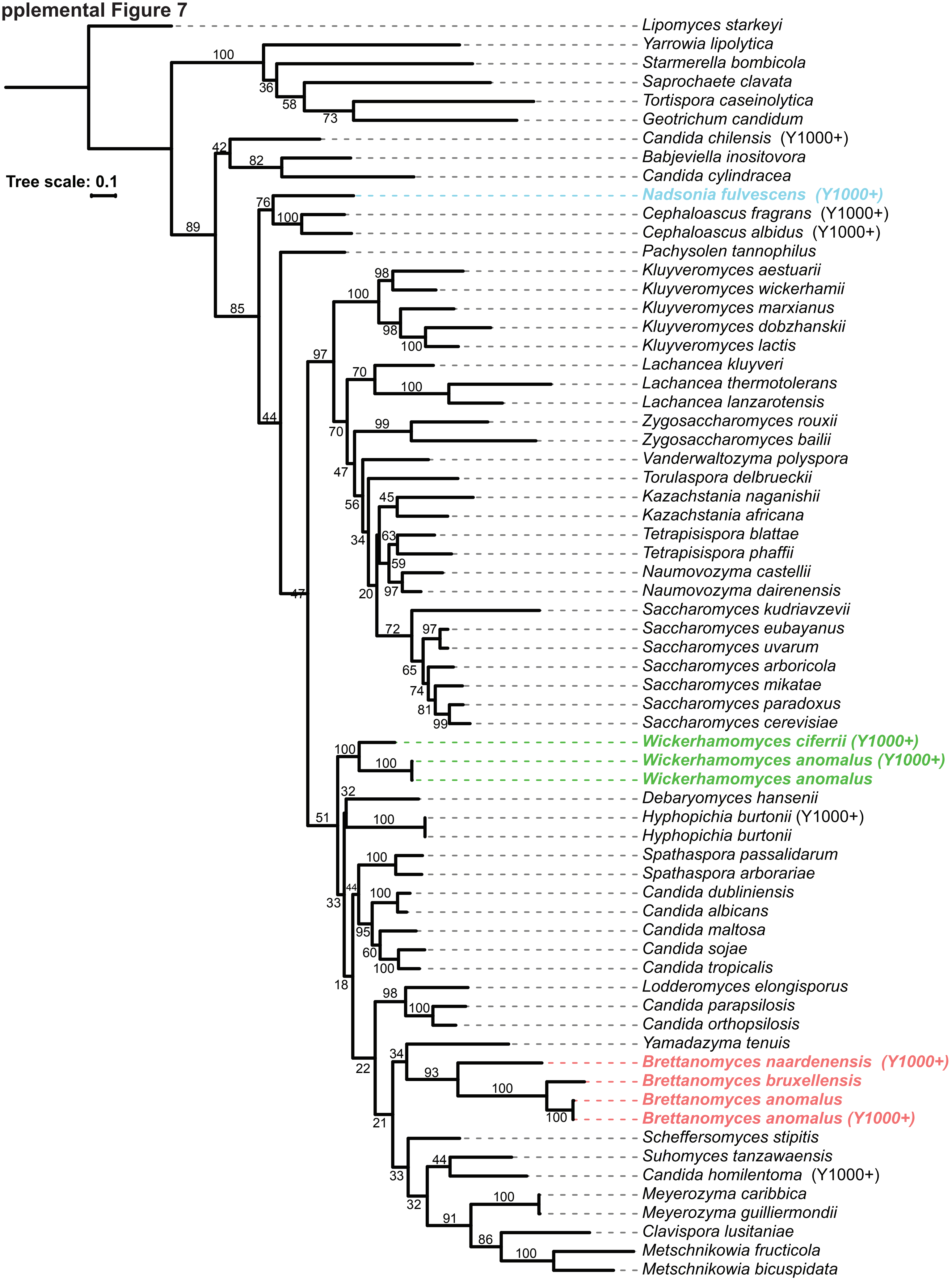

#### Supplemental Figure 8

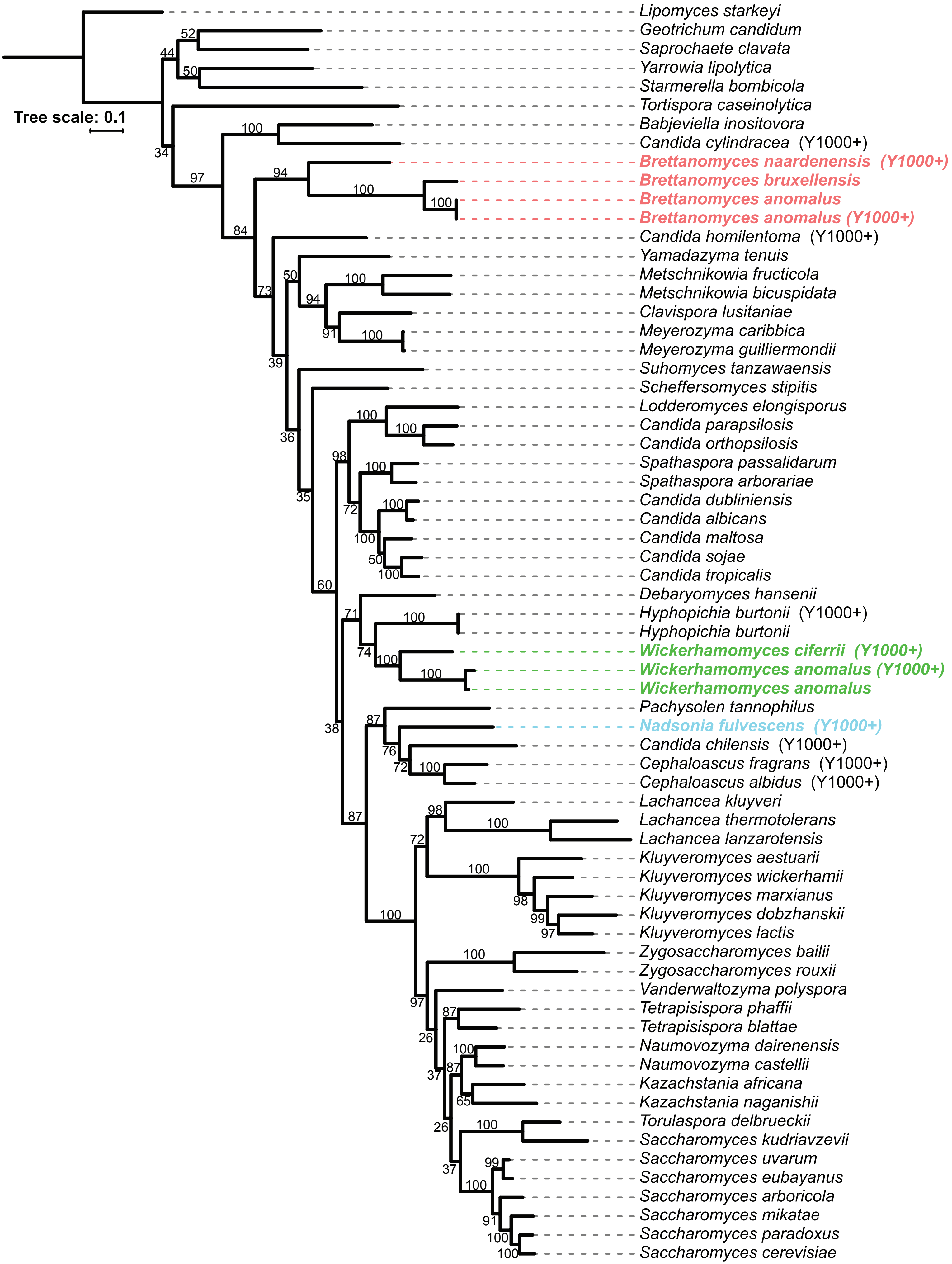

#### Supplemental Figure 9

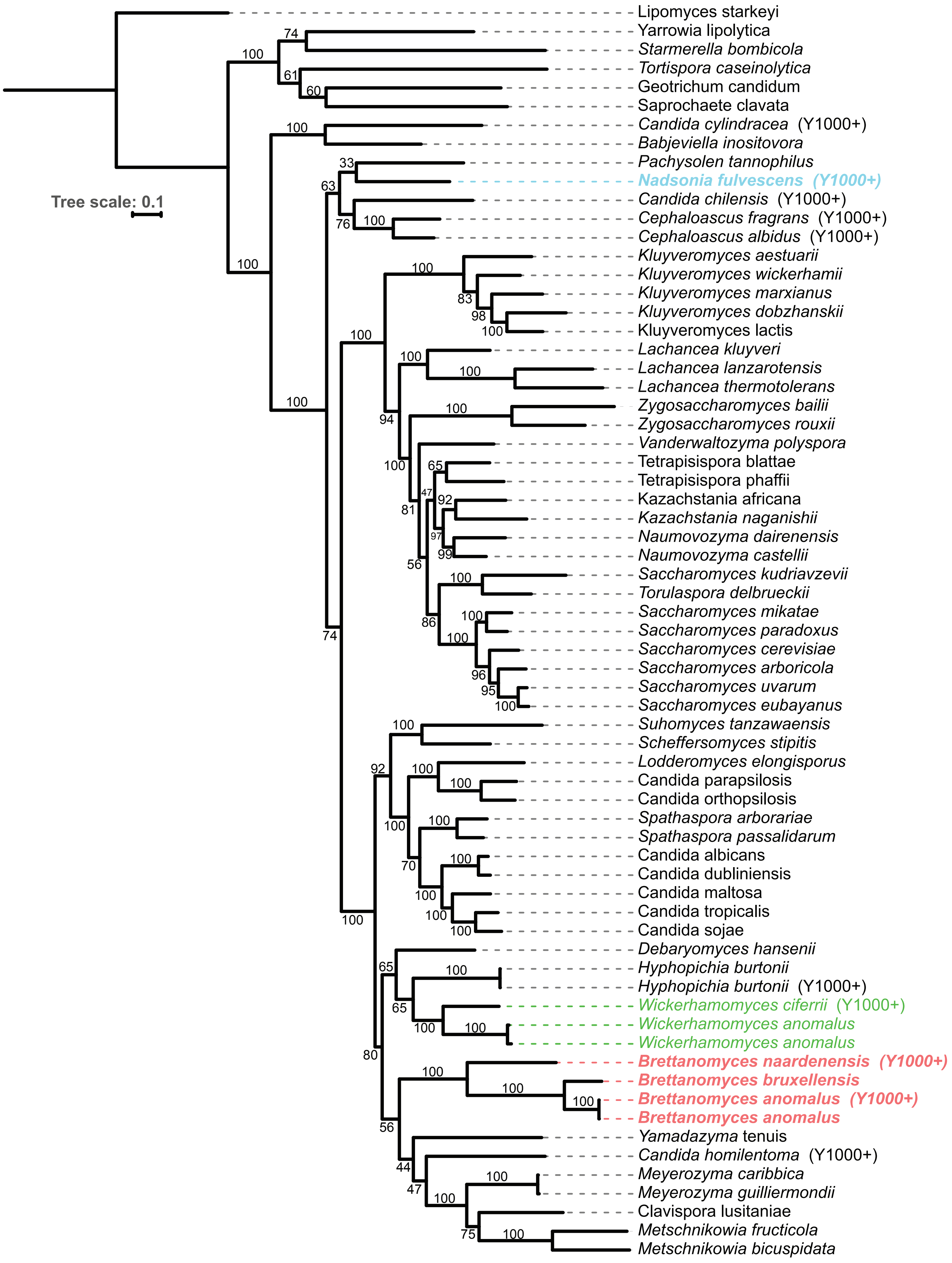

**Supplemental Figure 10**

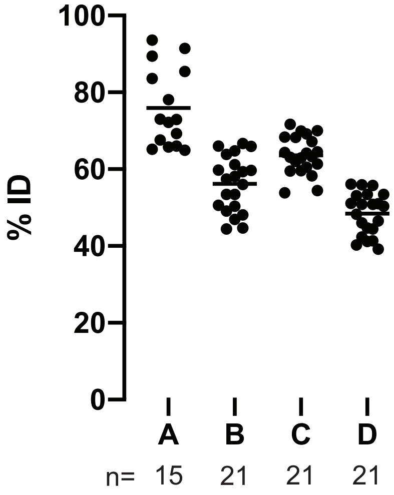
